# Electrodiffusion active pump model with asymmetric immersed chemical potentials

**DOI:** 10.64898/2025.11.30.691367

**Authors:** Pilhwa Lee

## Abstract

Electrodiffusion is essential in understanding the mechanisms of electrophysiology. Active exchange pumps are critical regarding volume homeostasis, and are also significant in the mechanisms of cell division, growth, and apoptosis. In the formalism of the immersed boundary (IB) method, we replace classical interface conditions across the membrane with regularized chemical potentials to control the permeation of each ionic species governed by the Poisson-Nernst-Planck equation. In asymmetry, to regulate ionic transport, continuous chemical potential barriers are augmented with energetic gradients represented by smoothed Heaviside kernels specifying the directions of active pumps. We obtain steady-state concentrations from electrodiffusion active pumps using Na^**+**^, K^**+**^, Cl^**−**^ ionic species, with background charges in the entire unified domain under periodic boundary conditions. As a consequence of the model simulation, electroneutrality, except for the thin space charge layers along the membrane, is well satisfied. The electrodiffusion active pump model for the exchange of sodium and potassium (NKE) exhibits a good fit to the theoretical formula over a broad range of perturbations in ionic concentrations, ensuring volume conservation in the steady state only when active pumps are functioning. It is also shown that van’t Hoff’s law is satisfied without active pumps. This is a foundation for applying the IB electrodiffusion active pump model for subcellular transport of water and molecules, possibly involving cell motility and migration.

## 1 Introduction

Brownian ratchet, a fundamental mechanism of actin-based cell motility and migration, with asymmetric potential walls represents active chemical potential with ATP hydrolysis [1, 2]. As one specific mode of microscopic machineries of directed transport with ATP hydrolysis, for example, rotary protein motors utilize chemical potentials with ATPase [3, 4]. In this article, we introduce asymmetric chemical potentials to represent passive and active transport of ionic species in the superposition of regularized singular integrals of Dirac delta and Heaviside kernels.

Cell membranes are water-permeable, but many macromolecules (e.g., sugar) are not permeable across the membrane [5]. By this water semi-permeable property of the membrane, we observe diverse osmotic effects of osmosis. For instance, the water transport in the epithelial cell [6], the osmotic power in MEMS [7], the DNA packaging in the vesicle [8], and the osmophoresis [9, 10] are all involved with them. Regarding volume homeostasis, in the presence of a non-uniform distribution of concentration between intracellular and extracellular domains and highly concentrated negatively charged macromolecules inside (e.g., proteins, nucleic acids), without elasticity of the plasma membrane and exchange pumps working together, the cell inevitably swells [11–13]; the volume swelling of a dead cell is a good example. Active exchange pumps involved in cellular volume regulation are also significant in the mechanism of cell division, growth, and apoptosis [14].

In electrodiffusion in subcellular scales, along the plasma membrane, the assumption of electroneutrality is not satisfied. There come the electrical double layers, so-called the *Debye layer* or *space charge layer*. In physiological ionic concentrations, the thickness of the space charge layers is around 10 nm. This is deeply involved in electro-osmosis and active ion pumps for cell migration [10, 15], even though these studies do not represent the space charge layers in finite dimensions. Electrokinetic phenomena, the hydrodynamics of electrolytes driven by an electrical field, are all related to this mechanism, and they are significant in the flow or particle control in micro and nanofluidics. The related primary physical structure is again the *Debye layer* generated at the interface between fluid and dielectric or conducting structure as described by [16]. The accompanying migration of a structure immersed in fluids is called electrophoresis. The electrophoresis has been applied to protein and molecule separation by capillary electrophoresis, as shown in [17, 18], and diagnostics described in [19]. In the context of proteomics on a chip, the details are referred to [20, 21]. The biological cell also implements active pumping mechanisms for spontaneously generated electric fields and induced migration [22]. The concentration gradient generated from external electric fields and space charge layers induces electrophoresis coupled with osmotic effects. The mobility of the vesicle is enhanced by more than 100% in the case of water semi-permeable vesicles [23].

Osmotic effects have been modeled with chemical potential barriers [24–26]. In the formalism of the immersed boundary method [27–29], which is mathematical and computational framework for fluid-structure interaction [30], chemical potential barriers are applied for the drift-diffusion of solutes or electrically charged ions, realizing their mediation of osmotic effects. Active pump models have been approached in macroscopic points of view [31–34], and specifically for cellular volume regulation and water transport [35–38]. Within the macroscopic pump-leak models, electrodiffusion was addressed by [39, 40], but prescribing electroneutrality uniformly even in the space charge layers. There is one approach where surface charge layers are represented in the electroneutral asymptotic limit of electrodiffusion with its well-posedness [41].

In experiments, surface charge layers were shown to be significant, involving phospholipids for closing the feedback loop in cellular protrusion, polarity, and migration [42]. Indeed, cell polarity is involved with Na^+^/H^+^ exchange pumps [43]. Most ion channels use surface charges to increase the rates of ionic entrance [44].

The present article focuses on the electrodiffusion active pump model with the immersed chemical potentials, in the formalism of the IB method. The prescribed immersed chemical potentials represent both passive permeation and active pumping. With the presence of regularized bell-shaped chemical potential barriers, passive ionic transport does exist. Concurrently, regularized Heaviside chemical potentials work as directed active pumps. Still, overall, there comes a non-equilibrium steady state with the proposed model. We have observed macroscopic van’t Hoff’s law from microscopic kinetic interaction among semi-permeable membranes and solutes via immersed chemical potentials in the sense of fluctuation-dissipation without active pumps. The cases of membranes as free boundaries floating with and without active pumps are mainly explored in the main article, followed by the cases of fixed boundaries. The proposed model does demonstrate active pumping essential for cellular volume conservation while *capturing the finite dimension of the space charge layers along the membrane*.

## 2 MATHERMATICAL FORMULATION

*D*_*i*_: diffusion coefficient of the *i*^th^ ionic species

*q*: the elementary electrical charge (charge on a proton)

*qz*_*i*_: charge of the *i*^th^ species

*K*_B_: Boltzmann constant

*T* : absolute temperature (degrees in Kelvin)

Ω_E_ : Eulerian domain

*ε*: dielectric constant

The notations for the variables are the following:

*x* : Eulerian coordinate

*X*_a_ and *X*_b_ : Lagrangian coordinates of the left and right membranes

*ψ*_*i*_(*x, t*): chemical potential of the *i*^th^ ionic species

Ψ_*w*_(*x*): chemical potential kernel with the influence range of 4*w*

*A*_*i*_(*t*): contribution of membrane to chemical potential of *i*^th^ ionic species

*c*_*i*_(*x, t*): concentration of the *i*^th^ ionic species

*F*_*i*_(*x, t*): flux per unit area of the *i*^th^ ionic species

*φ*(*x, t*): electrical potential

*ρ*_0_(*x*): background electrical charge density

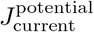: the current to potential map for the active pump

The left and right membranes are placed at the positions *X*_a_ and *X*_b_. These two membranes represent two free boundaries from the cross-section of the cell in the *x* direction.

### 2.1 The chemical potential

The chemical potential is expressed in the following way:

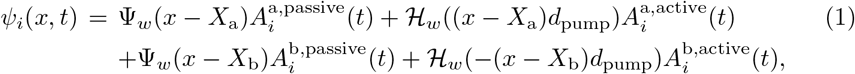

where the chemical potential amplitudes of 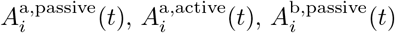, and 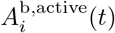 are generally defined as time-dependent, but in this article, set to be constant. They represent the contributions of passive and active transport of the membrane at *X*_a_ and *X*_b_ to the chemical potential barrier for the *i*^th^ species. The chemical potential kernel Ψ_*w*_ defines how the contribution 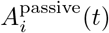 is to be spread out in space in the neighborhood of the membrane with the compact support of 4*w* [29]. Active pumps are prescribed with the smoothed Heaviside kernel *H*_*w*_ and 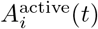 specifying the directionality of active pumping and the energetic gradients, respectively. When *d*_pump_ is 1, the active pump works outwardly. On the other hand, when *d*_pump_ is −1, the active pump works inwardly. In general, any bell-shaped function with compact support could be used for the chemical potential kernel Ψ_*w*_. Here, we make use of the smoothed Dirac delta function of the immersed boundary method [30], which is defined as follows:

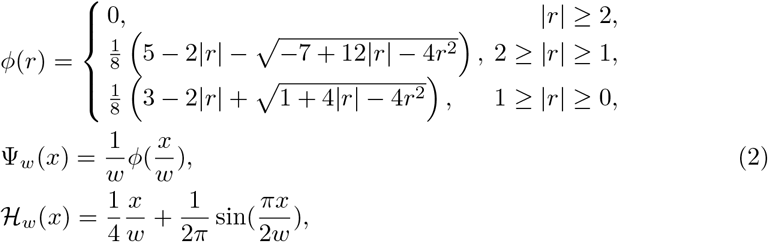

where *w* is a scaling factor such that Ψ_*w*_ has a support of width 4*w*. *H*_*w*_(*x*) has a transitional band width that is a function of *w, f* (*w*).

### 2.2 The electrostatic potential: the Poisson equation

The electrical potential is a solution of the Poisson equation:

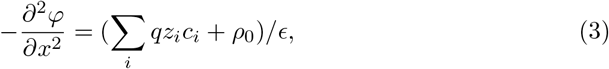

where *ρ*_0_ represents the background electrical charge density. The background charges reside on large molecules such as proteins or nucleic acids, and the net background charge is typically negative. The background charge density is prescribed to satisfy the local electroneutrality in the entire domain at *t* = 0. This equation is to be solved on a periodic domain. Because of this, it is necessary for the existence of a solution that the integral of the right-hand side over the domain is zero, i.e., that the system as a whole is electrically neutral (even though electroneutrality can be violated locally). In practice, electroneurality is also satisfied locally to a good approximation except in the space charge layer near the membrane (Figure 2(b) and (d)). Eq.(3) defines *φ* uniquely up to an additive constant. The choice of this constant has no significance since only the potential difference has physical effects.

For output purposes we define transmembrane voltages *V*_a_ and *V*_b_, as follows:

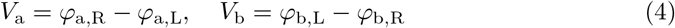

Where

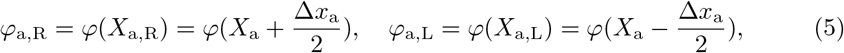

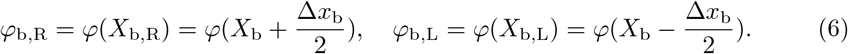

Here, Δ*x*_a_ is the distance between *X*_a,L_ and *X*_a,R_. In the same way, Δ*x*_b_ is the distance between *X*_b,L_ and *X*_b,R_. The computational electrode positions of *X*_a,L_, *X*_a,R_, *X*_b,L_, and *X*_b,R_ are chosen in a symmetric way with respect to the center line of the cell. They are outside the domains of the space charge layers and the support of the chemical potential, but close to the membranes. Their sign conventions are chosen so that they measure intracellular potential with respect to extracellular potential, where the space between the two membranes represents the intracellular space, and the rest of the domain represents the extracellular space.

### 2.3 The electrodiffusion equations

The electrodiffusion equations are formulated in the following way:

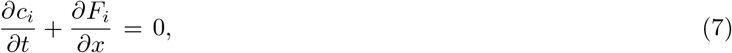

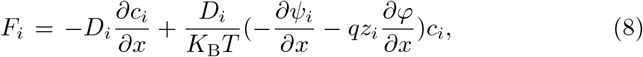

and the ionic currents from the electrodiffusion are the following:

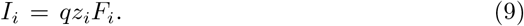

Eq.(7) is the conservation law (continuity equation) for the *i*^th^ species of ion. In this equation *c*_*i*_ is the concentration and *F*_*i*_ is the flux per unit area of this ionic species. Eq.(8) gives the flux per unit area as a sum of three terms: diffusion, drift induced by chemical potential, and drift induced by the electrical potential. In Eq.(9), *I*_*i*_ represents the current density (current per unit area) of *i*^th^ ionic species.

The Poisson equation and the electrodiffusion equations (Subsections 2.2 and 2.3) together constitute the Poisson-Nernst-Planck (PNP) system.

### 2.4 The chemical potentials for active and passive transports

The passive membrane permeation is represented by chemical potentials via a smoothed Dirac delta function (Figure 1(b), blue dotted profile). Active pumps are represented by chemical potentials with smoothed Heaviside functions (Figure 1(b), red dotted profile). The overall chemical potentials are constructed in a superposition of active and passive transmembrane transport (black solid profile).

**Fig. 1.**
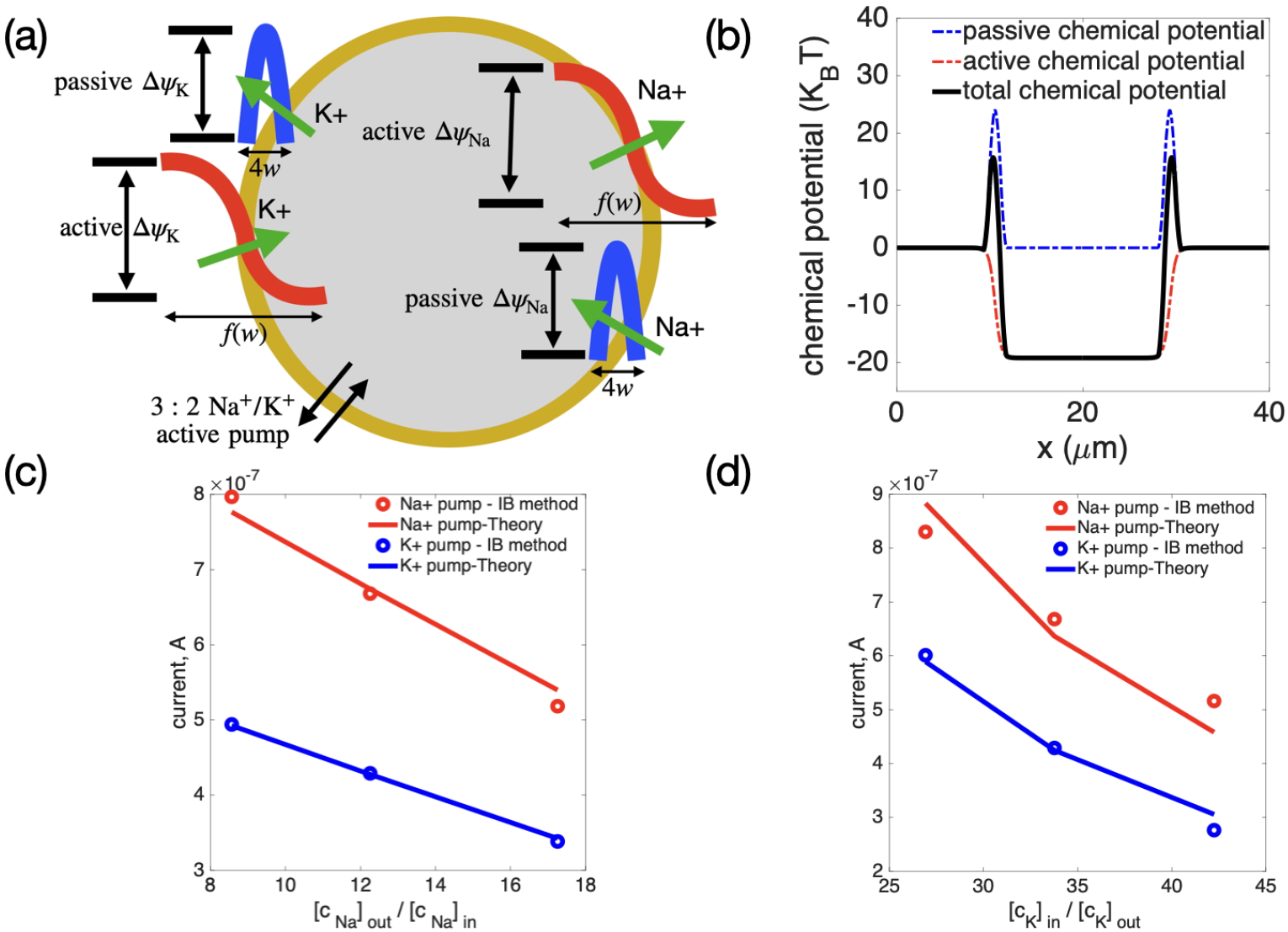
(a) Schematic representation of the active pump model with asymmetric chemical potentials. (b) Chemical potential distributions for active pumps (red dotted profile, smoothed Heaviside), passive membrane permeation (blue dotted profile, smoothed Dirac delta function), and superposition of active and passive transmembrane transport (black solid profile). (c) and (d) The fluxes of NKE pump, outflux of sodium ions (red circles) and influx of potassium ions (blue circles) computed from the immersed boundary (IB) method, are compared with the theoretical formula (Eq. 11), solid curves) with variation in the ratios of [Na^+^]_out_/[Na^+^]_in_ and [K^+^]_in_/[K^+^]_out_.

**Fig. 2.**
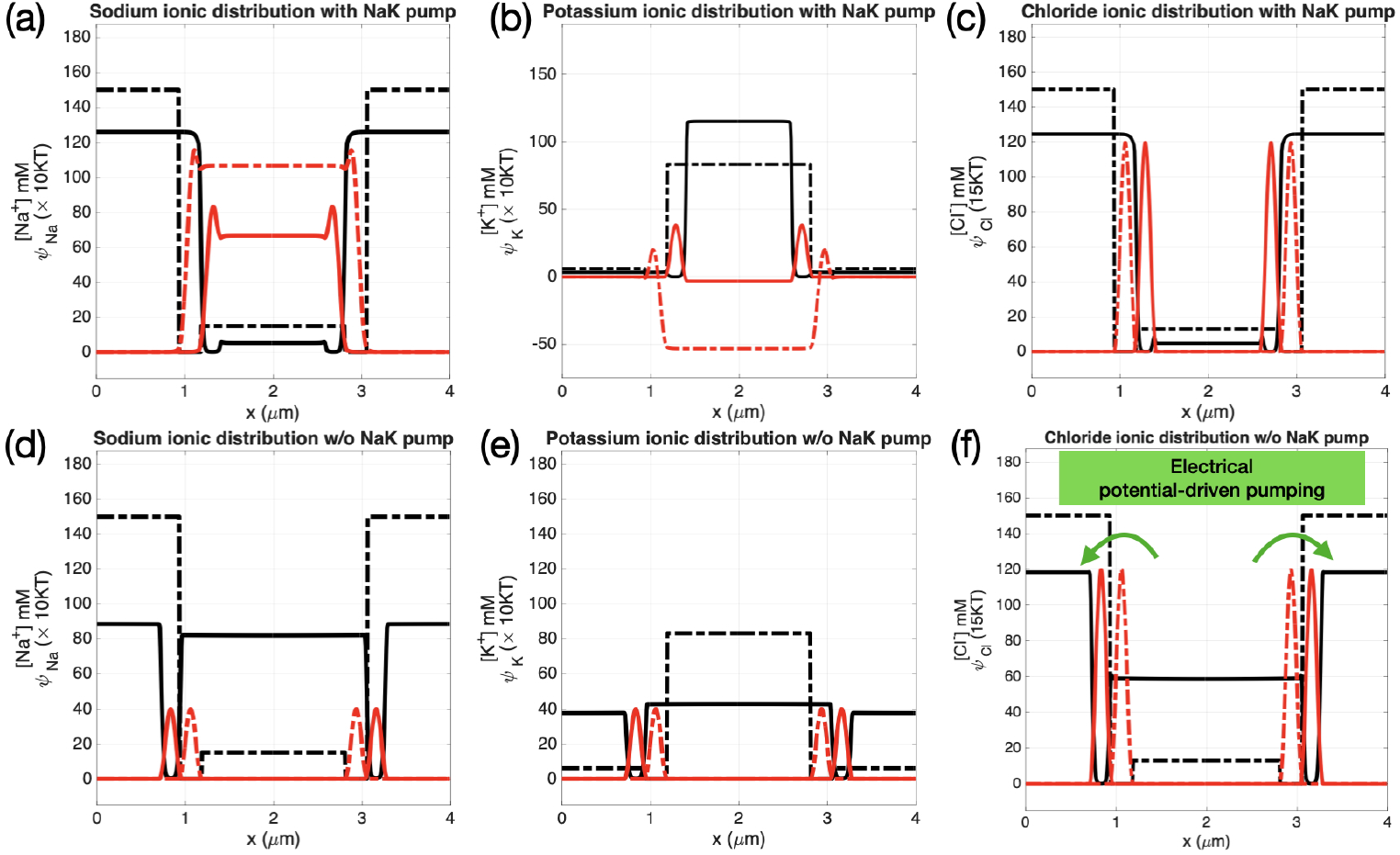
Free boundary: Ionic distributions associated with the chemical potentials. (a) Sodium ionic distribution with NaK pump, (b) Potassium ionic distribution with NaK pump, (c) Chloride ionic distribution with NaK pump, (d) Sodium ionic distribution without NaK pump, (e) Potassium ionic distribution without NaK pump, (f) Chloride ionic distribution without NaK pump. Solid and dotted black profiles are for the distributions at the steady state of *t*=120ms and at the initial *t*=0ms. Red profiles are for chemical potential distributions.

### 2.5 Active transport model formulation with current to potential map

In order to represent the active transport, the chemical potential is prescribed with its amplitude across the immersed boundaries in the following:

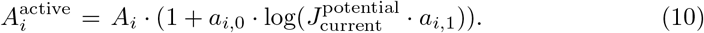

Here, the current-to-potential map, 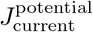does represent the main formulation of transmembrane current kernel in terms of ionic concentrations and membrane potentials applied to construct the chemical potential amplitudes 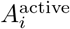 for active pumping of *i*^th^ ionic species.

### 2.6 NKE, Sodium-Potassium exchanger

The model is adapted from Li et al. [45]. The sodium and potassium currents are in the stoichiometry of 3:2 as counter currents.

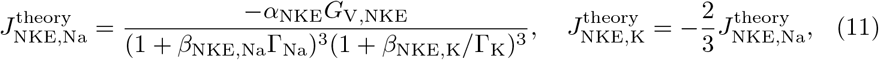

where 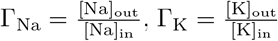, and 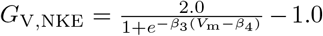. Accordingly, the current to potential map for the sodium-potassium active pump is prescribed in the following:

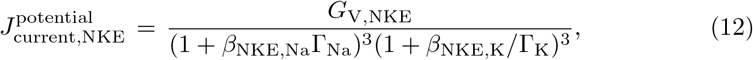

and the specific parameter values for the chemical potentials of active transport of sodium and potassium ionic species (Eq. 10) are in Appendix Table 2.

### 2.7 Free boundaries as chemical potential barriers interacting with elastic membranes

The one-dimensional membranes of *X*_*a*_ and *X*_*b*_ represent the cross-section of the cell. The cellular elastic tension at the membranes *X*_a_ and *X*_b_ is transmitted to the solutes/ions and fluid as *F*_ms_ and *F*_mf_, respectively in the sense of fluctuation and dissipation. For example, for the left membrane at the Lagrangian coordinate *X*_a_,

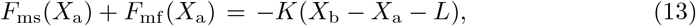

where *L* is the resting length at *t* = 0, and *K* is the elastic stiffness. Similarly, for the right membrane at the Lagrangian coordinate *X*_b_:

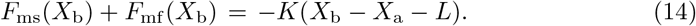

From the principle that the total integral of Eulerian chemical force densities from each ionic species with respect to *X* is the same as the Lagrangian force at *X*,

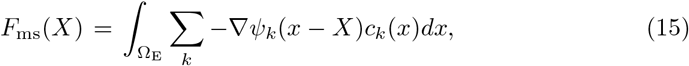

and *F*_mf_ can be explicitly formulated as follows:

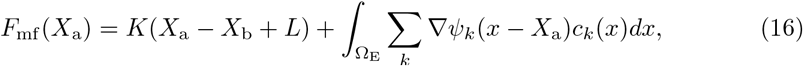

in other words, the elasticity of the cell membrane and chemical force are transduced to fluid from the kinetic collision between dissolved ionic species and the membrane.

In the Stokes flow in the 1D domain, the fluid flow is stationary (*u* = 0), and there remain only pressure gradients across the membranes (Figure 5). Accordingly, the relative sliding between the semi-permeable membrane and fluid, and the associated drag is induced by *F*_mf_ in the following:

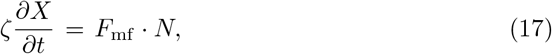

where *N* is the unit vector indicating the outer normal direction, and *ζ* is the osmotic stiffness. The distribution of hydrodynamic pressure can be obtained from the following stationary Stokes flow:

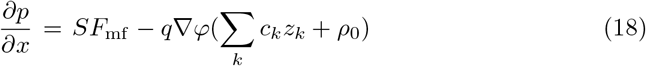

where the Lagrangian force from the membrane to the fluid, *F*_mf_ is spread out with the spreading operator *S* in the following:

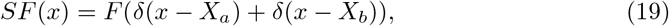

and the spontaneously generated electrical potential exerts an electrical force, so-called Lorentz force.

### 2.8 Solving the electrodiffusion equations of the Poisson-Nernst-Planck coupled to free boundaries

The computer simulation methodology of this paper is based on *the immersed boundary method with advection-electrodiffusion* [27, 28]; but here the membrane is fixed or floating in place, and fluid flows are stationary with only hydrodynamic pressure remaining. The entire domain is one-dimensional.

For the spatial domain of 5*µ*m, the spatial grids are 512 with Δ*x* = 9.766nm. The time-stepping is Δ*t* = 3ns. The main algorithmic flow for one time-stepping iteration is the following:

1. Apply Fast Fourier Transform (FFT) for solving the Poisson equation for the electrical potential, Eq. (3).
2. Construct chemical potentials for each ionic species, Eq. (2).
3. Incorporate a second-order Godunov-upwind method for reconstruction of concentration distributions for each ionic species [27].
4. Apply FFT for solving the hydrodynamic pressure distributions, Eq. (18).
5. Prescribe the advection of Lagrangian membrane coordinates of *X*_a_ and *X*_b_, Eq. (17).

The main numerical solvers of the partial differential equations is from the frameworks of PETSc [46].

### 2.9 Parameterization of chemical potentials

The parameters to be optimized for active chemical potentials are the following:

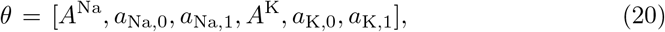

and PETSc Tao framework [46] with gradient-based optimization is incorporated to solve the following optimization problem:

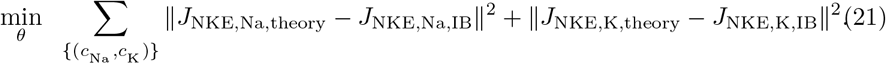

where the theoretical active currents of sodium and potassium, *J*_NKE,Na,theory_ and *J*_NKE,K,theory_ are from Eq. (11). *J*_NKE,Na,IB_ and *J*_NKE,K,IB_ are from the electrodiffusion active pump model currents with the immersed chemical potentials. Those currents are sampled at *X*_a_ and *X*_b_. The paired initial conditions of {(*c*_Na_, *c*_K_)} are from perturbations of sodium ionic concentration without change in potassium ionic concentration, and also from perturbations of potassium ionic concentration without change in sodium ionic concentration with the baseline concentration of Table 1 (Fig. 1(c) and (d)).

**Table 1.** Initial concentrations of all ionic species and background charges.

| Ion species | exterior concentration | interior concentration |
| --- | --- | --- |
| $\text{Na}^+$ | 150mM | 15mM |
| $\text{K}^+$ | 5mM | 100mM |
| $\text{Cl}^-$ | 150mM | 13mM |
| $\text{X}^-$ | 5mM | 102mM |

**Table 2.** Main parameters for the IB electrodiffusion active pump models.

| Parameters | values |
| --- | --- |
| $A_{Na}^{passive}$ | $5.0 \times 10^{-7} \text{ K}_B T$ |
| $A_K^{passive}$ | $5.0 \times 10^{-7} \text{ K}_B T$ |
| $A_{Cl}^{passive}$ | $1.0 \times 10^{-6} \text{ K}_B T$ |
| $A_X^{passive}$ | $5.0 \times 10^{-6} \text{ K}_B T$ |
| $A_{Na}$ | $3.333 \times 10^{-7} \text{ K}_B T$ |
| $A_K$ | $1.666 \times 10^{-7} \text{ K}_B T$ |
| $a_{Na,0}$ | 0.2 |
| $a_{Na,1}$ | $1.0 \times 10^6$ |
| $a_{K,0}$ | 0.5 |
| $a_{K,1}$ | $1.0 \times 10^6$ |
| $w_{passive}$ | $8 \Delta x$ |
| $w_{active}$ | $8 \Delta x$ |
| $\Delta x_a$ | $40 \Delta x$ |
| $\Delta x_b$ | $40 \Delta x$ |

## 3 Results and Discussion

### 3.1 NKE model construction

The current to potential map for the sodium-potassium active pump is prescribed in Eq. (12) based on the theoretical formulation of active exchange fluxes, Eq. (11). The results with fixed boundaries from the IB method with the immersed chemical potentials (solid curves) are fitted with the theoretical formula of the flux terms of the Na^+^/K^+^ active pump (NKE) (Fig. 1(c) and (d)), i.e., outflux of sodium ions (red circles) and influx of potassium ions (blue circles) with variations in the ratios of Γ_Na_ = [Na^+^]_out_/[Na^+^]_in_ and Γ_K_ = [K^+^]_in_/[K^+^]_out_ in the initial configuration of ionic concentrations.

### 3.2 Ionic distributions associated with the chemical potentials

The electrodiffusion active pump model is simulated with free membranes. The ionic distributions are shown to be adjusted with the moving boundaries in Figure 2. For comparison, the active pump is turned off and a moderate amount of sodium and potassium ions are flooded in and out of the membranes. Figure 2 shows sodium ionic distribution with and without NKE (Panels (a) and (d)), potassium ionic distribution with and without NKE (Panels (b) and (e)), and chloride ionic distribution with and without NKE (Panels (c) and (f)). Solid and dotted black profiles are for the distribution at the steady state of *t*=120ms and at the initial *t*=0ms. Red profiles are for chemical potential distributions.

### 3.3 Electrical and chemical potential distributions and time courses

The electrical potentials and electrical charge densities are shown with free boundaries in Figure 3 with and without active pumps. Electrical and chemical potential distributions are shown with and without NKE (Panels (a) and (c)), electrical charge density distribution with and without NKE (Panels (b) and (d)). Solid curves and dotted curves are for *t*=120ms and *t*=6ms. Active components of the chemical potentials are represented by solid curves for sodium (blue) and potassium (red) ionic species. Passive components of the chemical potentials for sodium (black) and potassium (green) are represented by dotted curves (Panel (e)). Membrane potential time courses are shown with and without NKE (Panel (f)). Resting membrane potential is recapitulated around −80mV with the active pump. Without an active pump, the membrane is tonically depolarized unsteadily. The proposed electrodiffusion model does not prescribe local electroneutrality off the space charge layers, but is naturally well satisfied in the model simulation.

**Fig. 3.**
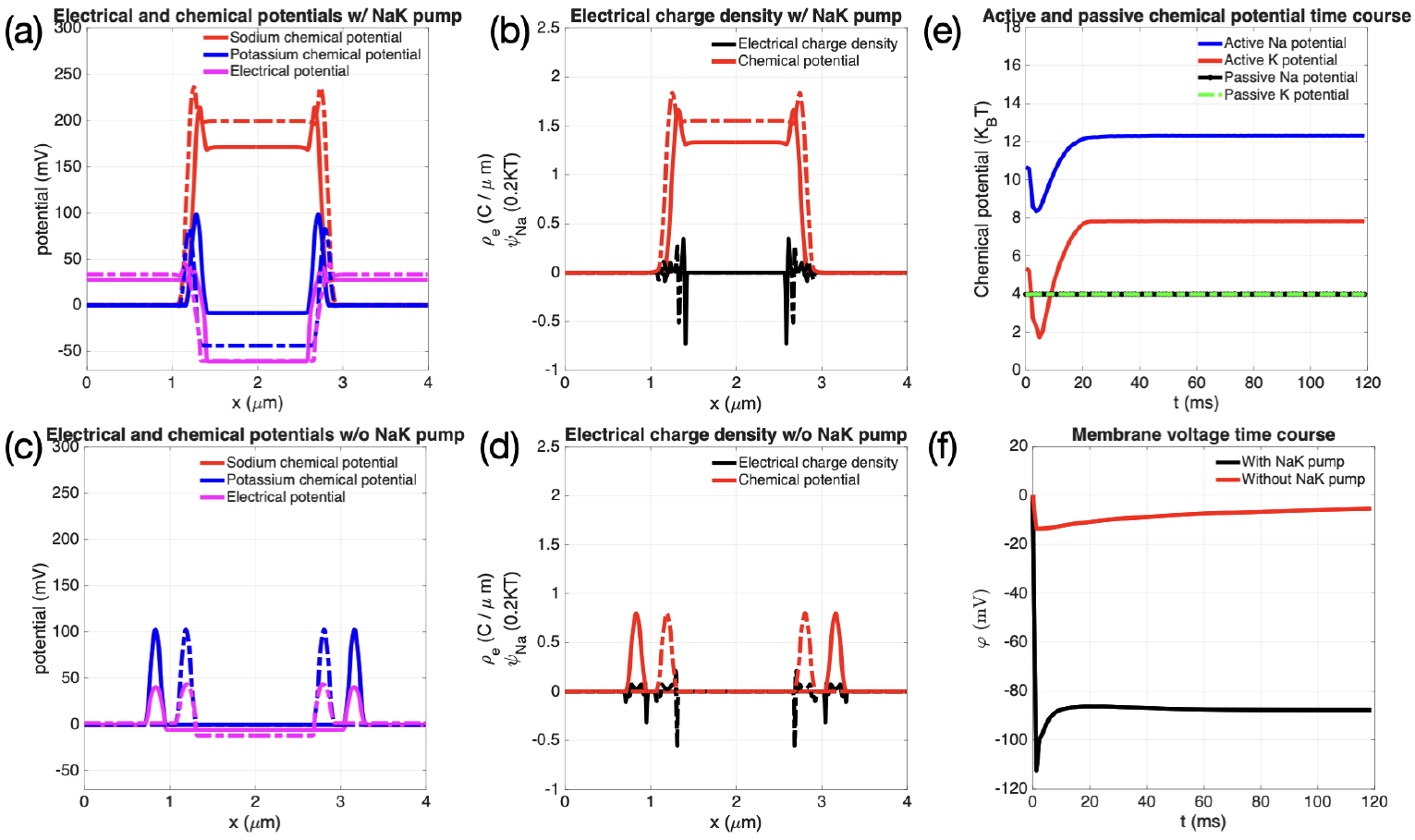
Electrical and chemical potential distributions and time courses. (a) and (c) Electrical and chemical potential distributions with and without the NKE pump. (b) and (d) Electrical charge density distribution with and without NKE pump. Solid curves and dotted curves are for *t*=120ms and *t*=6ms. (e) Active components of the chemical potentials for sodium (blue) and potassium (red) ionic species. Passive components of the chemical potentials for sodium (black) and potassium (green) are represented by dotted curves. (f) Membrane potential time courses with and without NKE pump.

### 3.4 Ionic current distributions and time courses

The time course of ionic currents (Na^+^, K^+^, Cl^−^, and total) as well as ionic current distributions is shown in Figure 4. The time courses of transmembrane ionic currents and total current with and without NKE in Panels (a) and (d). Ionic current distributions with and without NKE at *t*=5.0ms (Panels (b) and (e)) and at *t*=19.2ms (Panels (c) and (f)). Chemical potentials for sodium and potassium ionic species are also shown in red solid and dotted profiles.

**Fig. 4.**
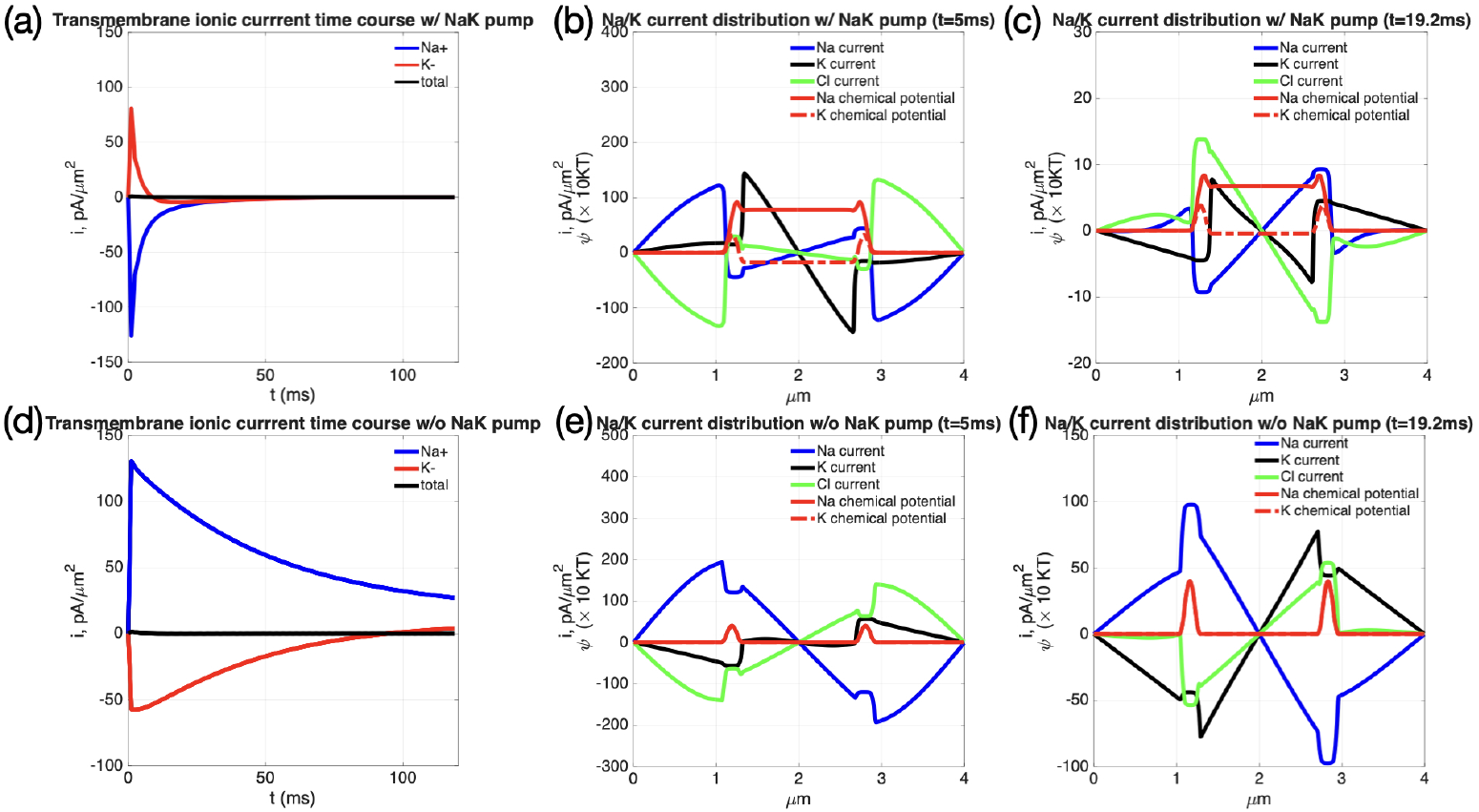
Ionic current distributions and time courses. (a) and (d) Transmembrane ionic currents and total current time courses with and without NKE pump. (b) and (e) Ionic current distributions with and without NKE pump at t=5.0ms. (c) and (f) Ionic current distributions with and without NKE pump at t=19.2ms. Chemical potentials for sodium and potassium ionic species are also shown in red solid and dotted profiles.

**Fig. 5.**
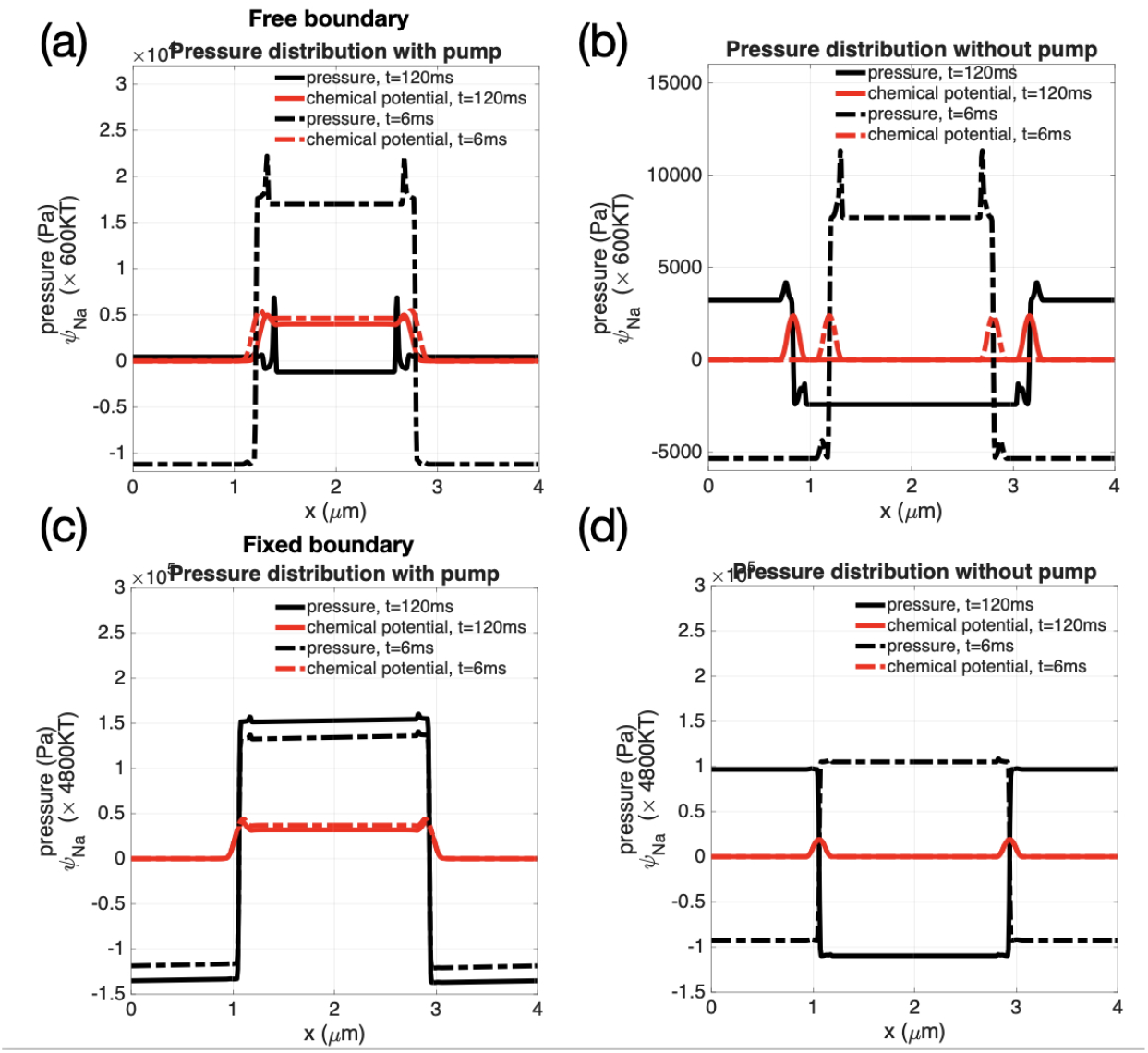
Pressure and checmial potential profiles. (a) and (b) Free boundary. Hydrodynamic pressure profiles with and without NaK pump. (c) and (d) Fixed boundary. Hydrodynamic pressure profiles with and without NaK pump. The solid and dotted profiles represent the time at *t*=120ms and *t*=6ms.

With the free boundaries, the ionic current distributions are more complex. In case the boundaries are fixed, the sodium and potassium currents across the boundaries are solely outward and inward, respectively (Fig. 9). On the other hand, when the boundaries are free, the sodium ionic currents are inward in the extracellular domain but outward in the intracellular domain. The case of potassium is more complicated. When the boundaries are free, the potassium ionic currents are inward in both the extracellular and intracellular domains at *t*=5ms. However, the potassium ionic currents turn out to be inward in the extracellular domain but outward in the intracellular domain at *t*=19.2ms. Chloride ionic currents are adjusted to satisfy the total current to be almost zero in the sense of electroneutrality.

When the active pump is turned off, sodium and potassium ionic currents are persistently inward and outward, respectively, i.e., passive permeation of those ionic species as the only transport across the membrane. Interestingly, the chloride ionic species has reverse flow in contrast to the case of working active pump, showing the involved pumping functionality implicitly without any explicitly prescribed active pumps in the way of total current to be almost zero in the sense of electroneutrality (Fig. 4(f)).

### 3.5 van’t Hoff’s Law in the microscopic representation of the interaction between the membrane and dissolved solutes in fluid

Hydrodynamic pressure profiles are shown in Figure 6. There is a dramatic change in hydrodynamic pressure across the intracellular to extracellular domains. The hydrodynamic pressure difference between the intracellular and extracellular domains is Δhydrodynamic pressure, and the osmotic pressure difference between the intra-cellular and extracellular domains is Δosmotic pressure. Figure 6(a) and (b) show Δhydrodynamic pressure versus Δosmotic pressure across the membranes as time courses of pressure and osmolarity differences with and without NKE. With active pumps working, Δhydrodynamic pressure is less than Δosmotic pressure, and the difference is through the work done by active pumps. Without active pumps working, Δhydrodynamic pressure is approximately equal to Δosmotic pressure, and this is consistent with the macroscopic law of van’t Hoff’s Law:

**Fig. 6.**
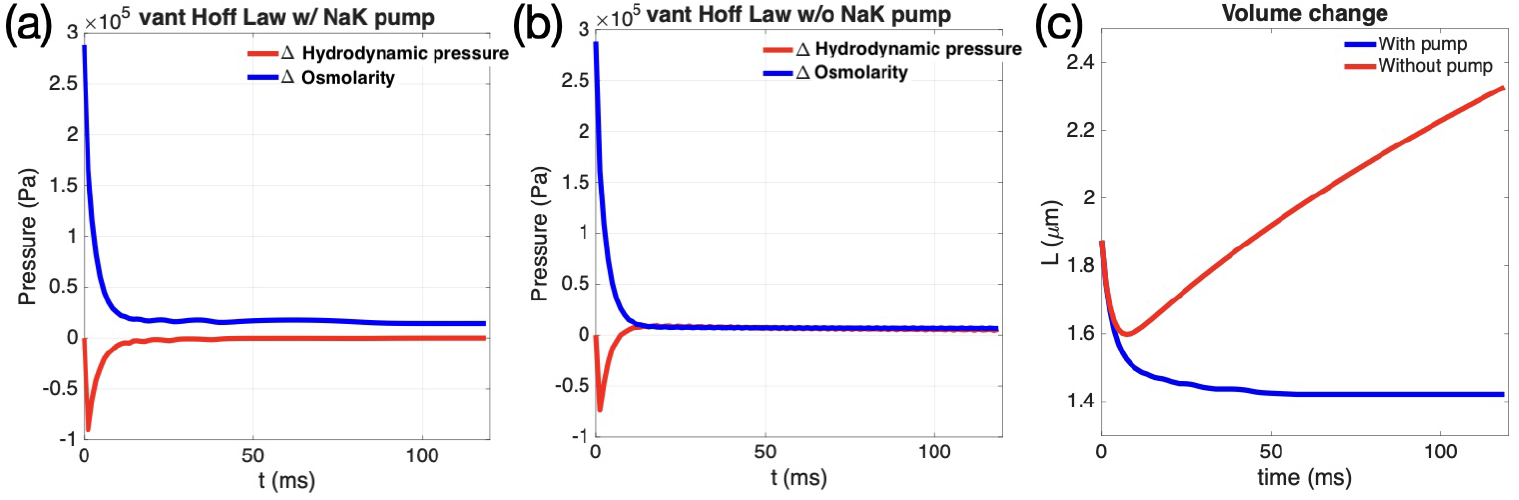
Hydrodynamic pressure versus osmotic pressure. (a) and (b) Time course of pressure and osmolarity differences across the membrane with and without the NKE pump. (c) Volume change with and without the NKE pump.

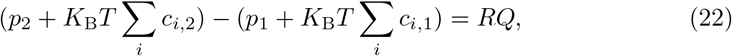

where *p*_1_ and *c*_*i*,1_ are the extracelullar hydrodynamic pressure and the concentration of *i*^th^ ionic species, and *p*_2_ and *c*_*i*,2_ are the intracellular hydrodynamic pressure and the concentration of *i*^th^ ionic species. All of them are out of the space charge layers. *R* is the water resistance, and *Q* is the water flux from the intracellular to extracellular domains, which is zero in the 1D model. Active pumps are essentially necessary for volume regulation. As shown in Fig. 6(c), the volume is preserved due to the active pumps after a transient adjustment in the beginning. On the other hand, the volume is immensely expanded to significant swelling when the active pump does not work.

### 3.6 The case of fixed boundaries

For the case of fixed boundaries, the electrodiffusion active pump model is simulated with membranes fixed. The ionic distributions are shown in Figure 7. For comparison, the active pump is turned off, and a moderate amount of sodium and potassium ions are flooded in and out of the membranes. The electrical potentials, electrical charge densities, and hydrodynamic pressures are shown in Figures 8 and 5 with and without active pumps. Resting membrane potential around −80mV is recapitulated with the active pump. Without an active pump, the membrane is depolarized. The electrodiffusion model does not prescribe electroneutrality outside the space charge layers, but is naturally well satisfied from model simulation. The time course of ionic currents (Na^+^, K^+^, Cl^−^, and total) as well as ionic current distributions is shown in Figure 9. When the active pump is working, as shown in Figure 9(a), the sodium current is outward, and the potassium current is inward. They gradually monotonically decrease with the adjusted ionic concentrations and electrical potentials. In the current active pump model, the sodium current is preserved to be outward at *t*=3ms and *t*=15ms, but the potassium current is switching from inward (*t*=3ms) to outward (*t*=15ms). Chloride ionic currents are adjusted to satisfy the total current to be almost zero in the sense of electroneutrality, except for the space charge layers.

**Fig. 7.**
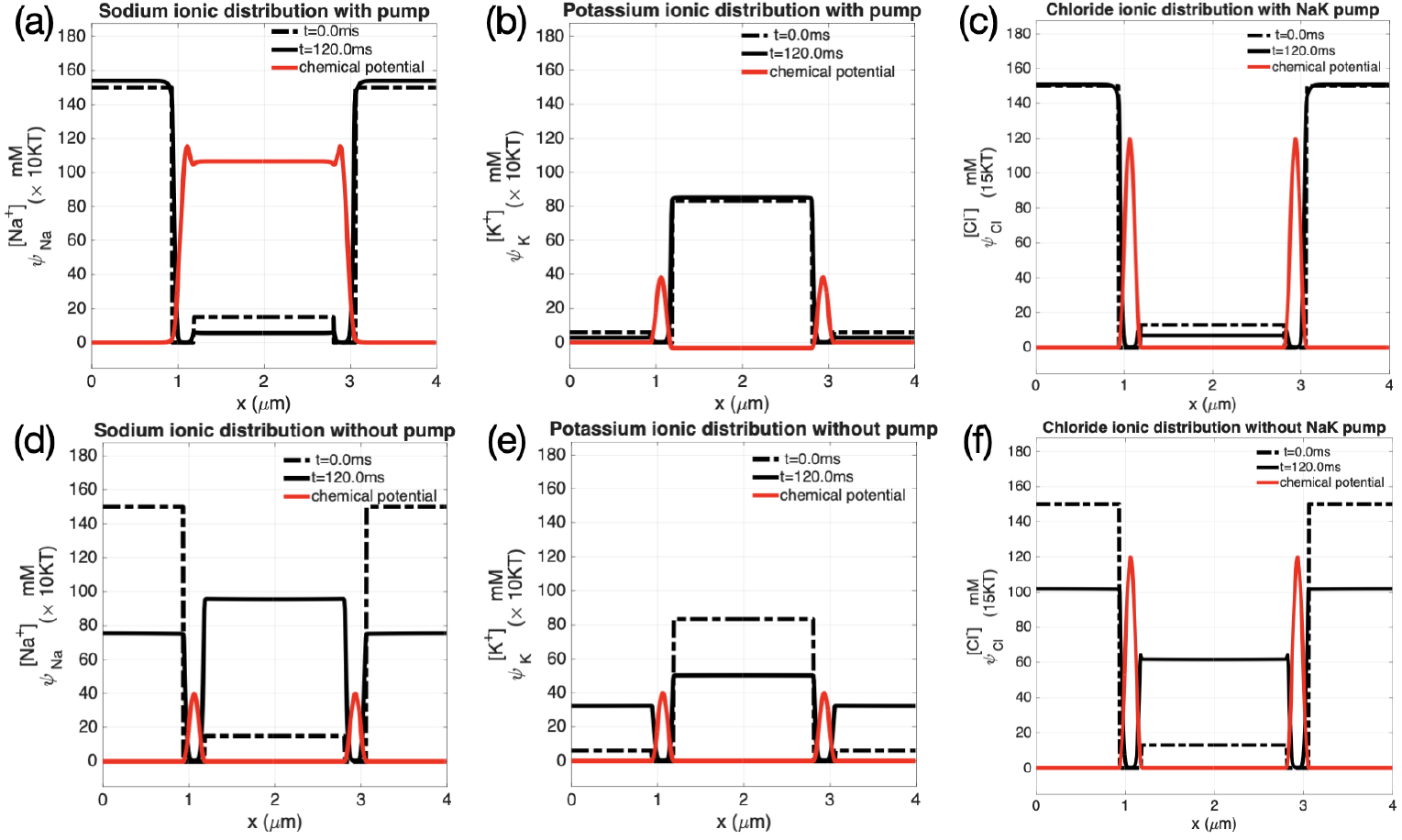
Fixed boundary: Ionic concentration distributions. (a) and (d) Sodium ionic distributions with and without NaK pump, (b) and (e) Potassium ionic distributions with and without NaK pump, (c) and (f) Chloride ionic distributions with and without NaK pump. Solid and dotted black profiles are for the distribution at t=120ms and t=0ms. Red profiles are for chemical potential distributions.

**Fig. 8.**
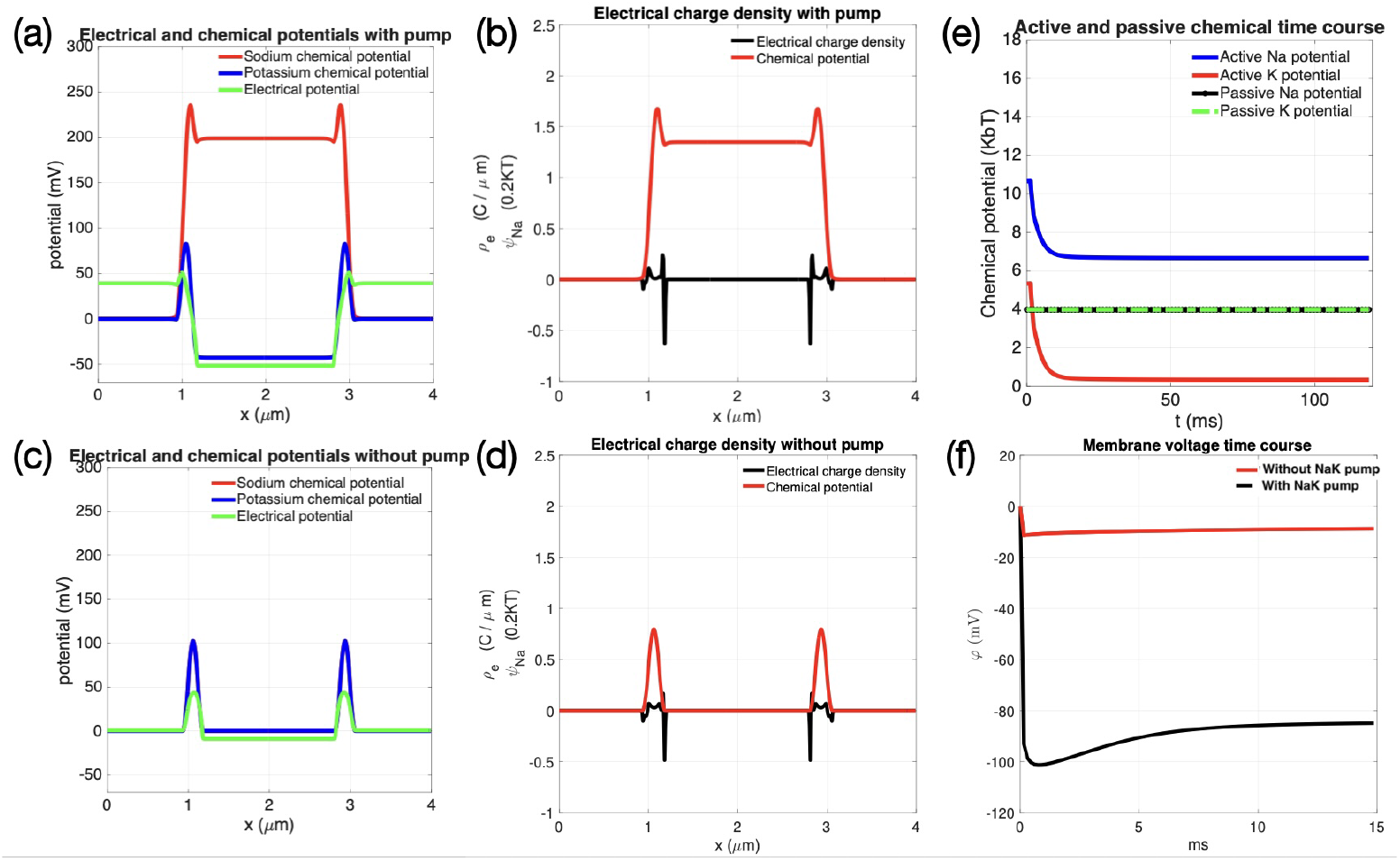
Fixed boundary: electrical and chemical potential profiles. (a) and (c) Electrical and chemical potential distributions with and without NaK pump, (b) and (d) Electrical charge density distributions with and without NaK pump. (e) Active components of the chemical potentials for sodium (blue) and potassium (red) ionic species. Passive components of the chemical potentials for sodium (black) and potassium (green) are represented by dotted curves. (f) Membrane potential time courses with and without NaK pump.

**Fig. 9.**
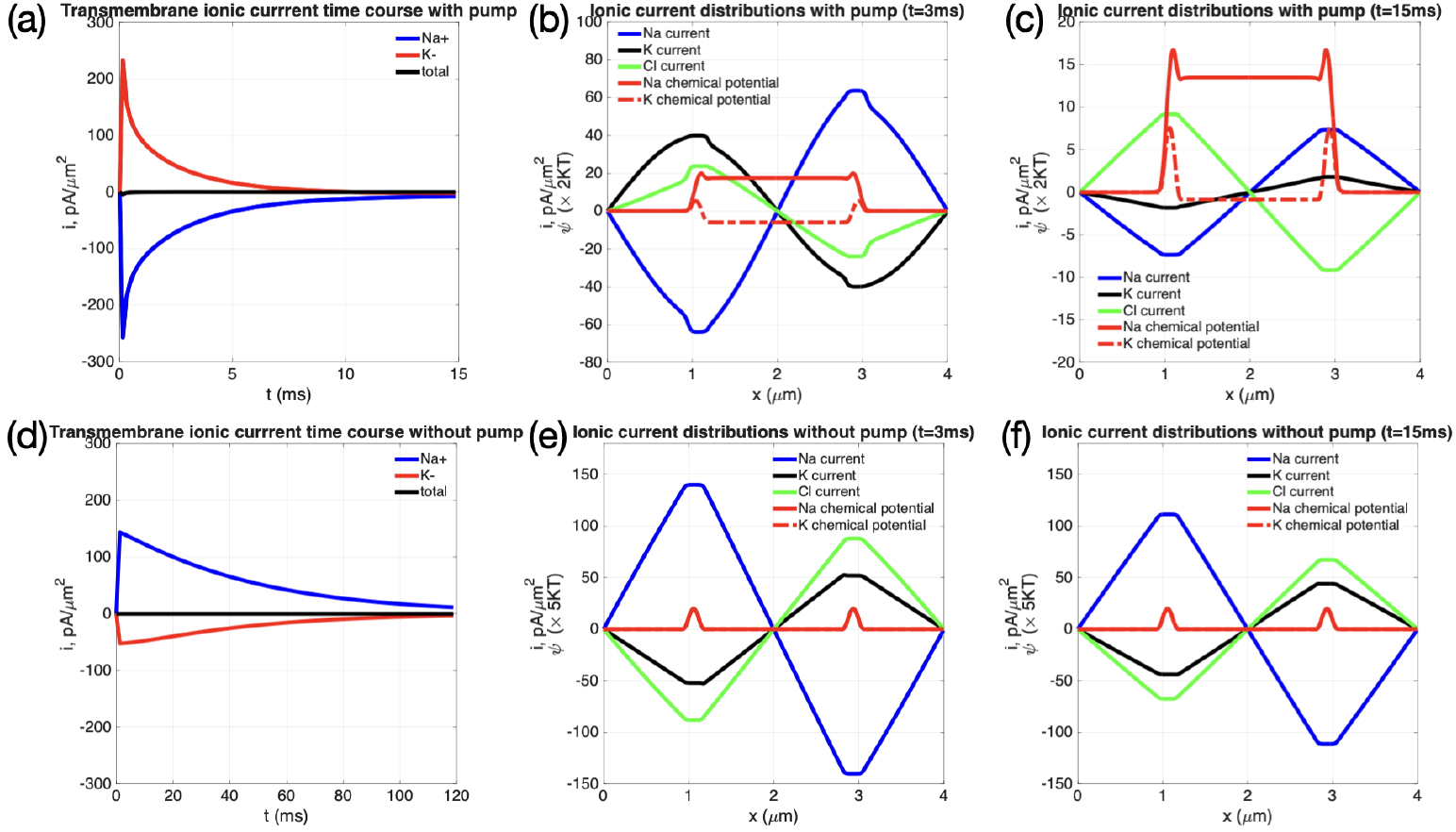
Fixed boundary: ionic current distributions and time courses. (a) and (d) Trans-membrane Ionic currents and total current time courses with and without NaK pump. (b) and (e) Ionic current distributions with and without NaK pump at t=3.0ms. (c) and (f) Ionic current distributions with and without NaK pump at t=15ms. Chemical potentials for sodium and potassium ionic species are also shown in red solid and dotted profiles.

When the active pump is turned off, sodium and potassium ionic currents are persistently inward and outward, i.e., passive permeation of those ionic species. Interestingly, the chloride ionic species has a reverse flow in contrast to the case of working active pump, practically pumping functionality is realized for chloride ions in the way of total current to be almost zero in the sense of electroneutrality, except for the space charge layers.

### 3.7 Membrane thickness

The prescribed membrane thickness is around 300 nm. In the involved discretization of the numerical methods, refining the mesh and grids around the membrane will make the domain of influence from the prescribed chemical potentials thinner, still in the non-asymptotic finite measure. Nevertheless, the competence of recapitulating physiological subcellular profiles of excitability in electrodiffusion is thought to be well conserved.

### 3.8 Generalizability for other types of active pumps

The proposed IB electrodiffusion active pump model focused on NKE is applicable to NHE (Na^+^*/*H^+^) and NKCC (Na^+^/K^+^/2Cl^−^) cotransporters. One conjecture is that the asymmetry in the superposition of regularized singular integral and Heaviside kernels is sufficient for arbitrarily prescribed active transport exchange currents in terms of intracellular and extracellular ionic concentrations and membrane voltage.

## 4 Conclusions

The newly developed IB electrodiffusion active pump model does reconstruct passive transport in the membranes via smoothed Dirac delta functions and induced currents from active pumps via smoothed Heaviside kernels. This is a new model of active pumps in the presence of the space charge layers in the electrodiffusion. The free boundaries without an active pump show van’t Hoff Law, justifying the microscopic representation of fluctuation-dissipation from membrane to fluid mediated by solutes/ions in the presence of chemical potentials. The model simulation reconstructs active pumping with volume preserved and swelling without active pumps while resolving the space charge layers in a non-asymptotic manner. An extension to 3D model simulations based on the IB electrodiffusion model of Poisson-Nernst-Planck equations coupled to Stokes flows remains for the future, where it will be intriguing to address the symmetry of Onsager’s law from asymmetric chemical potentials. This is a foundation for applying the immersed boundary electrodiffusion active pump model for subcellular transport of water and molecules, possibly involving cell motility and migration, such as dendritic spine motility and synaptic plasticity, and cellular migration, as well as cellular contraction, excitation-contraction coupling, and renal solute concentration.

## 5 Acknowledgements

This article is a celebration of Charlie Peskin’s 80th birthday. The author was partially supported by RCMI (U54MD013376) and ASCEND (UL1GM118973/ RL5GM118972) in the summer sabbatical research in Sean Sun’s lab at Johns Hopkins.

## Appendix

We consider three bio-ions, each with a different concentration between the two membranes from its concentration in the regions outside the two membranes. The initial concentrations are shown in Table 1. The symbol *X*^−^ in the table denotes the background charges.

Data will be made available on reasonable request.

## References

[1] Peskin, C.S., Odell, G.M., Oster, G.F.: Cellular motions and thermal fluctuations: 16 the Brownian ratchet. Biophysical Journal 65, 316–324 (1993)

[2] Peskin, C.S., Ermentrout, B., Oster, G.: The correlation ratchet: a novel mechanism for generating directed motion by ATP hydrolysis. In: V.C. Mow, e.a. (ed.) Cell Mechanics and Cellular Engineering. Springer, New York (1994)

[3] Oster, G., Wang, H.: Rotary protein motors. TRENDS in Cell Biology 13, 114–121 (2003)

[4] Sun, S.X., Wang, H., Oster, G.: Asymmetry in the F1-ATPase and its implications for the rotational cycle. Biophysical Journal 86, 1373–1384 (2004)

[5] Finkelstein, A.: Water Movement Through Lipid Bilayers, Pores, and Plasma Membranes : Theory and Reality. Distinguished Lecture Series of the Society of General Physiologists. Wiley, New York (1987)

[6] Spring, K.R.: Epithelial Fluid Transport–A Century of Investigation. News Physiol Sci 14, 92–98 (1999)

[7] Su, Y.-C., Lin, L., Pisano, A.P.: A Water-Powered Osmotic Microactuator. Journal of Microelectromechanical Systems 11, 6681–6692 (2002)

[8] Evilevitch, A., Lavelle, L., Knobler, C.M., Raspaud, E., Gelbart, W.M.: Osmotic pressure inhibition of DNA ejection from phage. Proc Natl Acad Sci USA 100, 9292–9295 (2003)

[9] Zinemanas, D., A. Nir: Osmophoretic Motion of Deformable Particles. International Journal of Multiphase Flow 21, 787–800 (1995)

[10] J Nardi, R.B., Sackmann, E.: Vesicles as Osmotic Motors. Physical Review Letters 82, 5168–5171 (1999)

[11] Dai, J., Sheetz, M.P., Wan, X., Morris, C.E.: Membrane tension in swelling and shrinking molluscan neurons. J Neurosci 18, 6681–92 (1998)

[12] O’Neill, W.C.: Physiological signicance of volume-regulatory transporters. American Journal of Physiology 276, 995–1011 (1999)

[13] Baumgarten, C.M.: Cell volume regulation in cardiac myocytes: a leaky boat gets a new bilge pump. J Gen Physiol 128, 487–489 (2006)

[14] Panayiotidis, M.I., Bortner, C.D., Cidlowski, J.A.: On the mechanism of ionic regulation of apoptosis: would the Na+/K+-ATPase please stand up? Acta Physiol (Oxf) 187, 205–215 (2006)

[15] Stroka, K.M., Jiang, H., Chen, S.-H., Tong, Z., Wirtz, D., Sun, S.X., Konstantopoulos, K.: Water permeation drives tumor cell migration in confined microenvironments. Cell 157, 611–623 (2014)

[16] Dukhin, S.S.: Non-equilibrium electric surface phenomena. Adv. Colloid Interface Sci. 44, 1–134 (1993)

[17] Hayes, M.A., Ewing, A.G.: Electroosmotic flow control and monitoring with an applied radial voltage for capillary zone electrophoresis. Anal Chem 64, 512–516 (1992)

[18] Lee, C.S., Blanchard, W.C., Wu, C.-T.: Direct control of the electroosmosis in capillary zone electrophoresis by using an external electric field. Anal. Chem. 62, 1550–1552 (1990)

[19] Landers, J.P.: Molecular diagnostics on electrophoretic microchips. Anal. Chem. 75, 2919–2927 (2003)

[20] Dolnik, V., Hutterer, K.M.: Capillary electrophoresis of proteins. Electrophoresis 22, 4163–78 (2001)

[21] Figeys, D.P.: Proteomics on a chip : Promosing developments. Electrophoresis 22, 208–216 (2001)

[22] Lammert, P.E., Prost, J., Bruinsma, R.: Ion drive for vesicles and cells. J. Theor. Biol. 178, 387–391 (1996)

[23] Fransaer, J., Celis, J.-P., Roos, J.R.: Electro-osmophoresis of a charged permeable microcapsule with thin double layer. J. Colloid Interface Sci. 151, 26–40 (1991)

[24] Atzberger, P.J., Kramer, P.R.: Theoretical framework for microscopic osmotic phenomena. Phys. Rev. E 75, 061125 (2007)

[25] Atzberger, P.J., Isaacson, S., Peskin, C.S.: A microfluidic pumping mechanism driven by non-equilibrium osmotic effects. Physica D 238, 1168–1179 (2009)

[26] Wu, C.-H., Fai, T.G., Atzberger, P.J., Peskin, C.S.: Simulation of osmotic swelling by the stochastic immersed boundary method. SIAM J. Sci. Comput. 37, 660–688 (2015)

[27] Lee, P.: The immersed boundary method with advection-electrodiffusion. PhD thesis, Courant Institute of Mathematical Sciences, New York University (2007)

[28] Lee, P., Griffith, B.E., Peskin, C.S.: The immersed boundary method with advection-electrodiffusion with implicit timestepping and local mesh refinement. Journal of Computational Physics 229, 5208–5227 (2010)

[29] Lee, P., Sobie, E., Peskin, C.S.: Computer simulation of voltage sensitive calcium ion channels in a dendritic spine. Journal of Theoretical Biology 11, 87–93 (2013)

[30] Peskin, C.S.: The Immersed Boundary Method. Acta Num. 11, 479–517 (2002)

[31] Finkelstein, A.: Carrier model for active transport of ions across a mosaic membrane. Biophysical Journal 4, 421–440 (1964)

[32] Bunow, B.: Chemical reactions and membranes: a macroscopic basis for faciliated transport, chemiosmosis and active transport part i: Linear analysis. J. Theor. Biol. 75, 51–78 (1978)

[33] Endresen, L.P., Hall, K., Hoye, J.S., Myrheim, J.: A theory for the membrane potential of living cells. European Biophysics Journal 29, 90–103 (2000)

[34] Sala, L., Mauri, A.G., Sacco, R., Messenio, D., Guidoboni, G., Harris, A.: A theoretical study of aqueous humor secretion based on a continuum model coupling electrochemical and fluid-dynamical transmembrane mechanisms. Communications in Applied Mathematics and Computational Science 14, 65–102 (2019)

[35] H J., Sx S.: Cellular pressure and volume regulation and implications for cell 20 mechanics. Biophys J 105, 609–619 (2013)

[36] J T., Sx S.: Active biochemical regulation of cell volume and a simple model of cell tension response. Biophys J 109, 1541–1550 (2015)

[37] Yellin, F., Li, Y., Sreenivasan, V.K.A., Farrell, B., Johny, M.B., Yue, D., Sun, S.X.: Electromechanics and volume dynamics in nonexcitable tissue cells. Biophys J 114, 2231–2242 (2018)

[38] Chodhury, M.I., Li, Y., Mistriotis, P., Vasconcelos, A.C.N., Dixon, E.E., Yang, J., Benson, M., Maity, D., Walker, R., Martin, L., Koroma, F., Qian, F., Konstantopoulos, K., Woodward, O.M., Sun, S.X.: Kidney epithelial cells are active mechano-biological fluid pumps. Nature Communications 13, 1–13 (2022)

[39] Mori, Y.: Mathematical properties of pump-leak models of cell volume control and electrolyte balance. J. Math. Biol. 65, 875–918 (2011)

[40] Jackle, J.: The resting potential of nerve cells and the Na,K-pump. arXiv:1703.03766 (2017)

[41] Stinchcombe, A.R., Mori, Y., Peskin, C.S.: Well-posed treatment of space-charge layers in the electroneutral limit of electrodiffusion. Communications on Pure and Applied Mathematics 69, 2221–2249 (2015)

[42] Banerjee, T., Biswas, D., Pal, D.S., Miao, Y., Iglesias, P.A., Devreotes, P.N.: Spatiotemporal dynamics of membrane surface charge regulates cell polarity and migration. Nature Cell Biology 24, 1499–1515 (2022)

[43] Patel, H., Barber, D.L.: A developmentally regulated Na-H exchanger in dictyostelium discoideum is necessary for cell polarity during chemotaxis. Journal of Cell Biology 169, 321–329 (2005)

[44] Green, W.N., Andersen, O.: Surface charges and ion channel function. Annu. Rev. Physiol. 53, 341–359 (1991)

[45] Li, Y., Zhou, X., Sun, S.X.: Hydrogen, bicarbonate, and their associated exchangers in cell volume regulation. Frontiers in Cell and Developmental Biology 9, 683686 (2021)

[46] Balay, S., Gropp, W.D., McInnes, L.C., Smith, B.F.: Efficient management of parallelism in object oriented numerical software efficient management of parallelism in object oriented numerical software libraries. In: Arge, E., Bruaset, A.M., Langtangen, H.P. (eds.) Modern Software Tools in Scientific Computing. Birkhauser Press, Basel (1997)

